# Cellular zinc status alters chromatin accessibility and binding of transcription factor p53 to genomic sites

**DOI:** 10.1101/2023.11.20.567954

**Authors:** Leah J. Damon, Daniel Ocampo, Lynn Sanford, Taylor Jones, Mary A. Allen, Robin D. Dowell, Amy E. Palmer

## Abstract

Zinc (Zn^2+^) is an essential metal required by approximately 2500 proteins. Nearly half of these proteins act on DNA, including > 850 human transcription factors, polymerases, DNA damage response factors, and proteins involved in chromatin architecture. How these proteins acquire their essential Zn^2+^ cofactor and whether they are sensitive to changes in the labile Zn^2+^ pool in cells remain open questions. Here, we examine how changes in the labile Zn^2+^ pool affect chromatin accessibility and transcription factor binding to DNA. We observed both increases and decreases in accessibility in different chromatin regions via ATAC-seq upon treating MCF10A cells with elevated Zn^2+^ or the Zn^2+^-specific chelator tris(2-pyridylmethyl)amine (TPA). Transcription factor enrichment analysis was used to correlate changes in chromatin accessibility with transcription factor motifs, revealing 477 transcription factor motifs that were differentially enriched upon Zn^2+^ perturbation. 186 of these transcription factor motifs were enriched in Zn^2+^ and depleted in TPA, and the majority correspond to Zn^2+^ finger transcription factors. We selected TP53 as a candidate to examine how changes in motif enrichment correlate with changes in transcription factor occupancy by ChIP-qPCR. Using publicly available ChIP-seq and nascent transcription datasets, we narrowed the 50,000+ ATAC-seq peaks to 2164 TP53 targets and subsequently selected 6 high-probability TP53 binding sites for testing. ChIP-qPCR revealed that for 5 of the 6 targets, TP53 binding correlates with the local accessibility determined by ATAC-seq. These results demonstrate that changes in labile zinc directly alter chromatin accessibility and transcription factor binding to DNA.

## Introduction

Regulation of gene expression is at the heart of virtually every biological process. How genes are switched on and off at a precise time and in a specific place drives developmental programs, defines the type and state of a cell, regulates cell fate decisions, and underlies basic physiology. The ability of transcription factors (TFs) to regulate gene expression is typically proportional to the accessibility of their binding sites in the genome.^1–3^ TFs either bind directly at promoters or in distal regulatory regions called enhancers, that are characterized by nucleosome depletion. DNase-seq^4^ and ChIP-seq^5,6^ have long been used to map regions of open chromatin and TF binding, respectively. TF binding can create a footprint of depletion of DNase-seq signal, although it has been reported that 80% of TFs don’t show detectable footprints.^7^ For TFs that do show footprints, DNase-seq has the advantage that it can capture TF-binding information for many TFs simultaneously. Consequently, several computational methods have been created to correlate DNase-seq signal with TF binding motifs.^8,9^ Because DNase-seq and ChIP-seq require millions of cells to generate high-quality datasets, an alternative approach for directly probing regions of open chromatin is becoming more widely used; the Assay for Transposase Accessible Chromatin with sequencing (ATAC-seq)^10,11^ requires only 500-50,000 cells. The transposase used in ATAC-seq inserts sequencing adapters into regions of open chromatin, and these regions correlate well with regulatory regions identified via DNase-seq and ChIP-seq.^10,12^ However, ATAC-seq does not allow for straightforward identification of which TFs may be active within these regions of open chromatin. Computational methods to identify TF “footprints” have been developed^13–15^ but these historically have not been as robust for ATAC-seq datasets as they are for DNase-seq datasets.^13^ Further, as previously noted, not all TFs leave detectable footprints and footprint depth is likely proportional to residence time on DNA.^7,16^

Transcription factor enrichment analysis (TFEA) is a recently developed computational approach that seeks to identify differentially active TFs in genomic datasets.^17^ TFEA takes regions of active transcription as input and subsequently uses DESeq2^18^ to establish a ranked list of differentially transcribed regions. Each of these regions is then subjected to TF motif scanning to determine which TF motifs are enriched in regions of open chromatin in a particular treatment, and an enrichment score (E-score) is calculated for each treatment relative to the control. TFEA was applied to nascent transcription datasets and accurately identified transcriptional responses to Nutlin-3a, lipopolysaccharide, and dexamethasone.^17^ TFEA can also be applied to ATAC-seq datasets to identify TF motifs associated with open chromatin and whether there is an increase or decrease in association in response to a given perturbation.^19,20^ Because regions of active transcription are associated with regions of open chromatin (both at enhancers and promoters), TFEA can thus be used to infer TFs that are enriched or depleted in genomic regions capable of active transcription in response to a given perturbation.

Of the more than 1600 known human TFs, approximately half (48%) bind zinc.^21^ While the specific mode of zinc ion (Zn^2+^) coordination can vary, many of these TFs use zinc-fingers to bind to DNA. Loss of Zn^2+^ results in a dysfunctional protein, with one of the most well annotated examples being TP53 and its tumorigenic R175H mutation.^22–24^ While this is evidence that some TFs may be susceptible to altered activities due to Zn^2+^ levels, historically the metal responsive transcription factor 1 (MTF1) is the only known TF whose activity is directly titratable by Zn^2+^ within the cell.^25–28^ However, we and others have shown that perturbation of cellular Zn^2+^ across multiple cell types results in global changes in transcription that are not directly related to MTF1.^29–31^ Although the concentrations of Zn^2+^ used in these studies were higher than expected physiological levels, even mild exposure to Zn^2+^ can alter the transcriptional landscape.^30^ Furthermore, it has been established that the labile Zn^2+^ pool of mammalian cells is dynamic, such that neuronal stimulation,^32^ cell-cycle progression,^33,34^ and fertilization,^35,36^ among other processes, can induce fluctuations to this labile pool in the pM to nM range. However, it is still unknown whether Zn^2+^ modulates transcription directly through its binding to transcription factors or indirectly through signaling pathways such as the MAPK pathway.^37^

In this study, we used ATAC-seq and TFEA to identify TFs that are putatively activated and/or repressed as a consequence of cellular Zn^2+^ status. Elevation of Zn^2+^ using ZnCl_2_ and depletion of Zn^2+^ using a Zn^2+^-specific chelator caused broad changes in the accessible chromatin landscape, even after a short 30 min perturbation. TFEA revealed 477 TF motifs that were differentially associated with newly open chromatin upon perturbation of Zn^2+^, where most motifs are reciprocally affected by Zn^2+^ and TPA. As one example, we found that the TP53 (p53) motif was globally enriched in regions of open chromatin upon Zn^2+^ depletion and reduced under Zn^2+^ replete conditions. Examination of how TFEA predictions translate to changes in TF binding revealed that for 5 of 6 selected TP53 target sites, p53 binding correlated with local accessibility, such that a decrease in accessibility led to decreased p53 binding and an increase in accessibility correlated with increased p53 binding. These results reveal that zinc status profoundly alters chromatin accessibility and this can be further propagated to zinc-dependent changes in TF binding to target genes, indicating that the functionality of zinc-dependent TFs can be sensitive to the labile Zn^2+^ pool.

## Results

To determine which TFs may be activated in Zn^2+^ replete or Zn^2+^ deficient conditions, we performed ATAC-seq on MCF10A cells subjected to perturbations of Zn^2+^. We first used a genetically encoded FRET sensor (NLS-ZapCV2)^38^ that is specific for measuring labile (freely exchangeable) Zn^2+^ in the nucleus. Addition of 30 µM ZnCl_2_ to the extracellular media caused a relatively slow but measurable rise in labile Zn^2+^ that peaks after 40 min (**Figure 1A**). To limit any secondary effects that may occur from downstream activation of TFs, we opted to collect cells for ATAC-seq after 30 min of ZnCl_2_ treatment. In situ calibration of the sensor combined with the binding parameters (K_D_’ = 5.3 nM, n = 0.29^30^) revealed that at 30 min there was a change in labile Zn from 24 ± 7.5 pM to 6.9 ± 2.9 nM. Addition of 50 µM of the Zn^2+^ chelator tris(2-pyridylmethyl)amine (TPA) resulted in a rapid decrease in nuclear Zn^2+^ to less than 1 pM (the minimum concentration of labile Zn^2+^ that can be quantified using NLS-ZapCV2). To keep treatments consistent, we also treated cells for 30 min with 50 µM TPA (**Figure 1B**).

**Figure 1.**
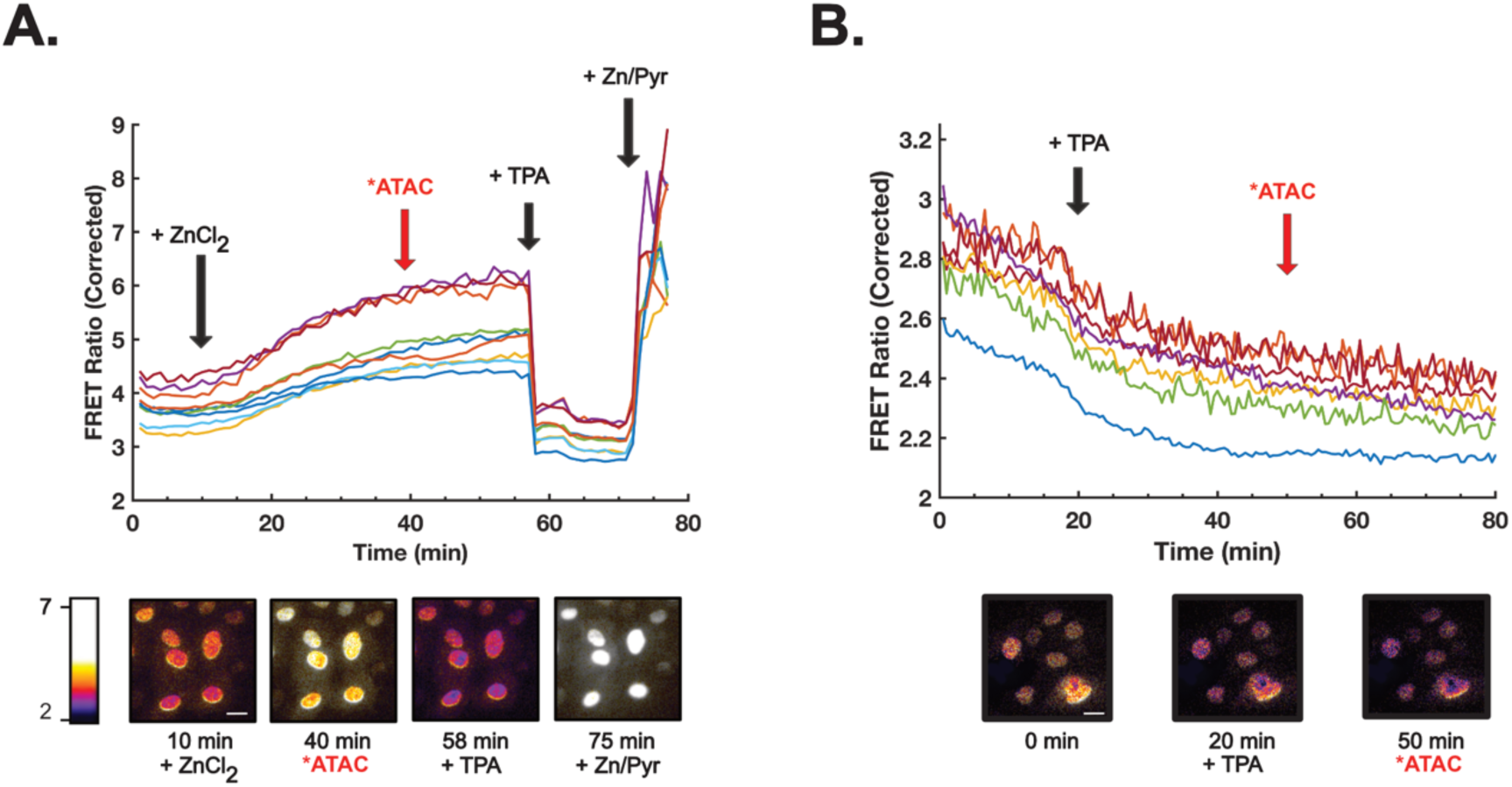
MCF10A cells are susceptible to perturbations in nuclear Zn^2+^. (A) Background-corrected FRET ratios for MCF10A cells expressing NLS-ZapCV2 (top) and representative cells at each of the indicated time points (bottom). Addition of ZnCl_2_ for 30 min results in an increase in labile Zn^2+^ from 24 ± 7.5 pM to 6.9 ± 2.9 nM. Addition of the Zn^2+^ chelator TPA at 58 min followed by the addition of Zn^2+^ and pyrithione at 75 min allows for calibration of the sensor and subsequent quantification of labile Zn^2+^. (B) Same as (A), but cells were treated with TPA at 20 min to deplete labile Zn^2+^. Each trace represents a single cell in the field of view. Lookup table values refer to the FRET ratio (background corrected FRET channel/background corrected CFP channel). Scale bar = 10 μm.

We performed ATAC-seq to assess which regions in the genome are differentially accessible in Zn^2+^ deficient and Zn^2+^ replete states. Treatment with TPA or ZnCl_2_ induced broad changes in global chromatin accessibility (**Figure 2A**). TPA treatment impacted accessibility more than ZnCl_2_ treatment (813 vs. 517 differentially accessible regions with TPA vs. ZnCl_2_) (**Figure 2B, Supplementary dataset 1**). In both treatments, there was an overall trend towards reduced accessibility, with 593 regions and 334 regions having negative log2FoldChanges, corresponding to 73% and 64% of the differentially accessible regions for TPA and ZnCl_2_, respectively. Interestingly, while treatment with TPA led to broad changes in accessibility, suggestive of non-specific global decreases in accessibility, elevated Zn^2+^ led to a smaller number of highly significant changes in accessibility in select regions, suggesting that perhaps elevation of Zn^2+^ leads to changes in specific regions of chromatin (**Figure 2A**). One of the top hits showing increased accessibility with ZnCl_2_ treatment was a region (chr16:56623012- 56628480; log2FC = 1.14, p_adj_ = 4.47E-07) directly aligned with the metallothionein isoform *MT1E*, a gene regulated by the metal regulatory transcription factor 1 (MTF1) in response to increased levels of Zn^2+^ **(Figure 3).** Additionally, the region overlapping a second metallothionein isoform, *MT2A*, also showed increased accessibility though to a lesser extent (chr16:56605751-56611835; log2FoldChange = 0.611, p_adj_ = 0.018). These results show that perturbations of the labile Zn^2+^ pool can alter the landscape of accessible chromatin in a short (30 min) time frame, with Zn^2+^ depletion leading to global decreases in chromatin accessibility and Zn^2+^ increases leading to more specific increases in accessibility. Further, we observe changes in accessibility in genomic regions corresponding to key zinc regulatory genes (*MT1E* and *MT2A*), as would be expected upon perturbation of metal homeostasis.

**Figure 2.**
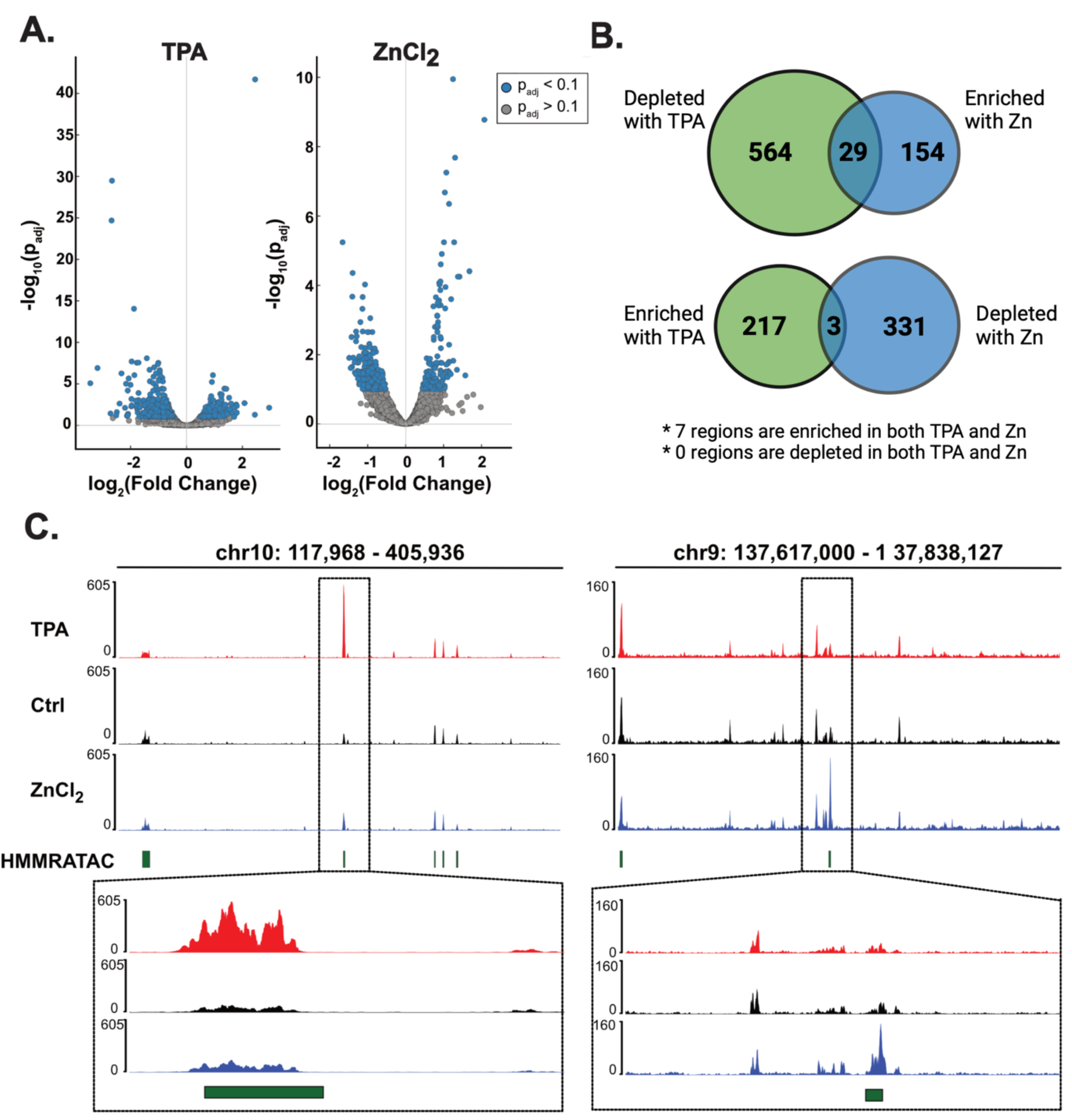
Perturbing cellular Zn^2+^ causes global changes in chromatin accessibility. (A) Volcano plots depicting genomic regions that exhibit a change in chromatin accessibility for cells treated with 50 µM TPA (left) or 30 µM ZnCl_2_ (right). A positive log2FoldChange indicates an increase in accessibility, while a negative log2FoldChange indicates a decrease in accessibility. (B) DESeq2 differential accessibility analysis shows that most peaks are uniquely accessible depending on Zn^2+^ status. The overlap in the Venn diagrams indicates the peaks which are inversely accessible between the denoted treatments. (C) Genomic tracks showing the top hit for differential accessibilities for TPA (left) and ZnCl_2_ (right) treated cells. Green boxes denote accessible chromatin as annotated using HMMRATAC.

**Figure 3.**
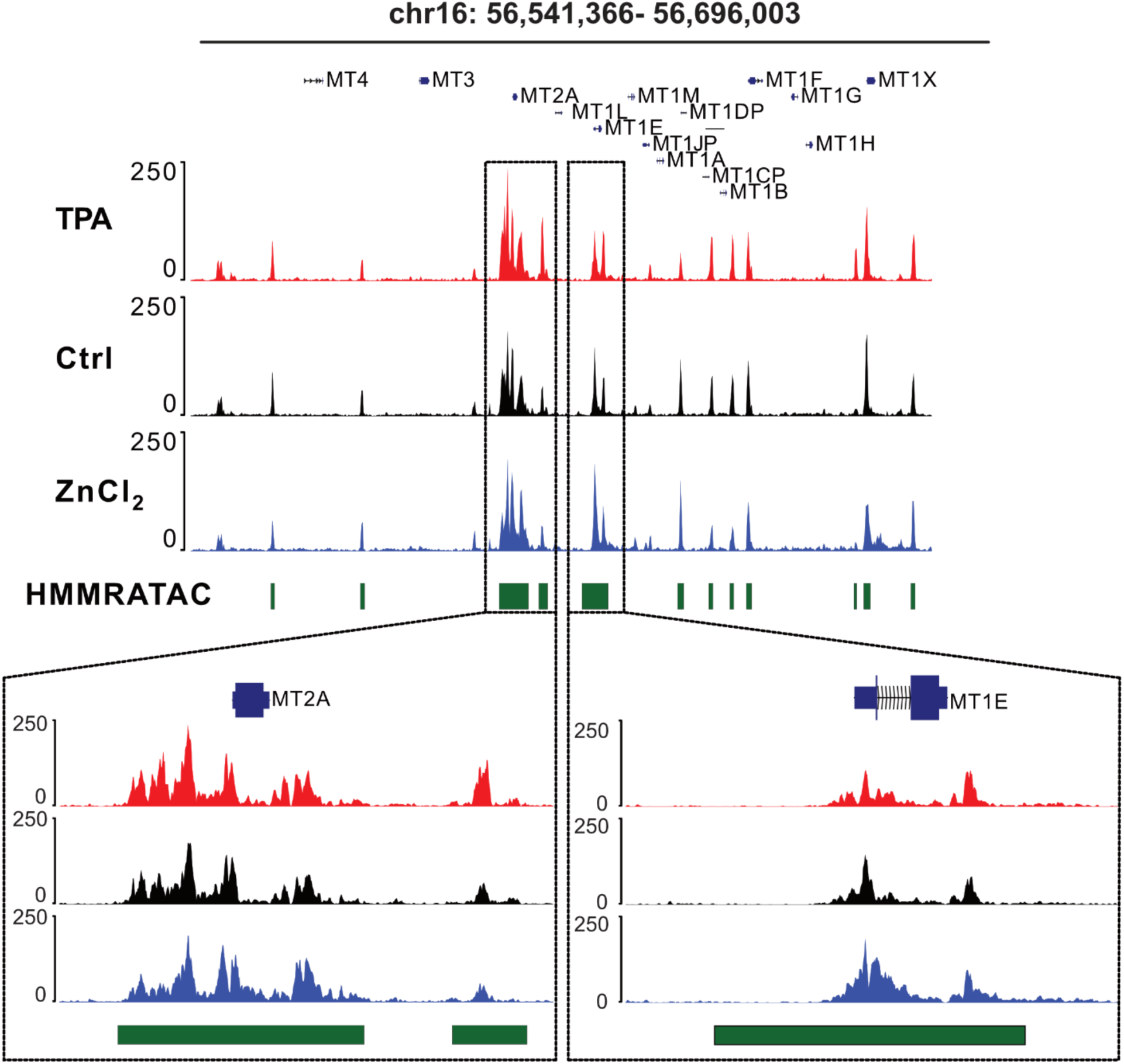
Genomic regions encompassing the Zn^2^ regulatory genes *MT2A* and *MT1E* show increased accessibility with ZnCl_2_ treatment. Top, the approximate 154 kb region of chromosome 16 where all MT isoforms reside. Bottom, zoomed in views of the *MT2A* locus (left) and the *MT1E* locus (right).

To determine which TFs may be activated or repressed in each treatment, we performed transcription factor enrichment analysis (TFEA).^17^ Briefly, regions of open chromatin were annotated as peaks using HMMRATAC^39^ and peaks from two technical replicates were combined using the muMerge algorithm.^17^ Then, each of 1279 TF motifs from HOCOMOCO^40^ were subjected to motif scanning using Find Individual Motif Occurrences (FIMO) to identify potential TF binding motifs within the detected peaks,^41^ and differential expression analysis using DESeq2 was performed to generate a ranked list of peaks that were differentially accessible between control and treatment conditions. This was used to generate an enrichment score (E-score) to determine which TF motifs were differentially enriched in regions of newly open chromatin between the control and treatment conditions. For all subsequent analyses, we removed motifs corresponding to TFs that are not transcribed in MCF10A cells as measured by PRO-seq (**Supplementary dataset 2)**. This reduced our TF motif list from 1279 motifs to 880 motifs, corresponding to 608 unique TFs.

Perturbation of Zn^2+^ led to broad changes in the enrichment of TF motifs associated with open chromatin. Elevation of Zn^2+^ to ∼ 7 nM resulted in significant (p_adj_ ≤ 1E-6) changes in enrichment of 449 TF motifs in regions of open chromatin, while reduction of Zn^2+^ to ∼ 1 pM using the chelator TPA resulted in changes in enrichment of 322 TF motifs (**Figure 4A, Supplementary dataset 3**). Strikingly, we saw a strong negative correlation between the E-scores of TPA and ZnCl_2_ (**Figure 4B**), suggesting reciprocal activation of these TFs depending on cellular Zn^2+^ status. Indeed, 38.9% (186/477) of differentially enriched motifs exhibited increased association with open chromatin in elevated Zn^2+^ and decreased association in TPA, while 22.6% (108/477) were depleted in high Zn^2+^ and enriched in TPA (**Figure 4C**). Further, a significant portion of motifs displayed altered enrichment upon Zn^2+^ elevation, with 17.6% (84/477) showing increases and 14.8% (71/477) showing decreases in regions of open chromatin. Conversely, there were very few motifs whose association with open chromatin was affected by TPA alone, of which 3.8% (18/477) were depleted and 2.1% (10/447) were enriched. The observation that elevation of Zn^2+^ induced changes in a substantial portion of the motifs (449/477, 94%) and on average these had higher E-scores than changes associated with TPA, suggests that acutely increasing the labile Zn^2+^ pool may have more significant effects on transcription compared to decreasing the labile pool.

**Figure 4.**
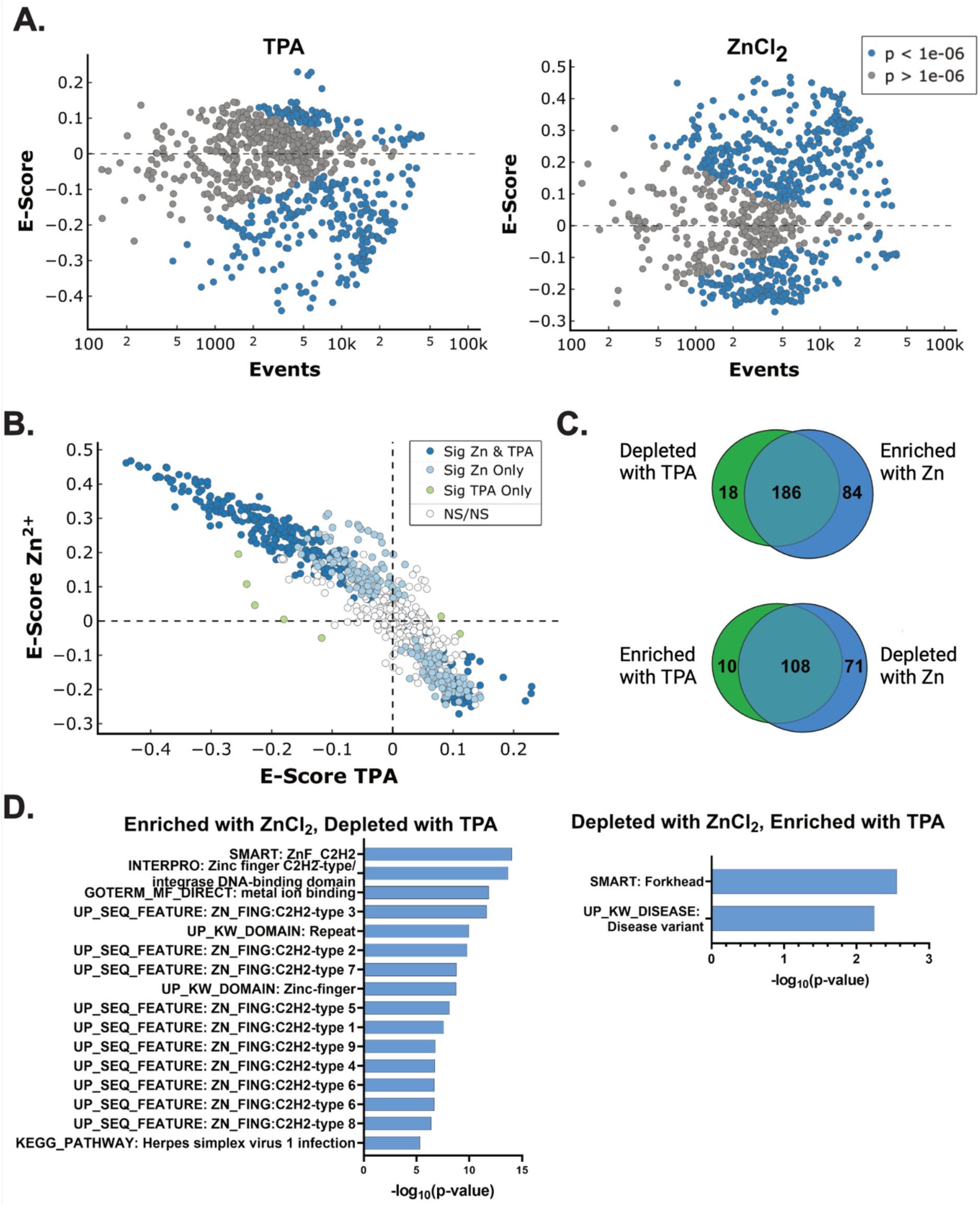
Perturbing cellular Zn^2+^ causes global changes in TF motif enrichment. (A) TFEA MA plots showing enrichment scores of 880 TF motifs. Cells were either treated with 50 µM TPA (left) or 30 µM ZnCl_2_ (right) for 30 mins, and enrichment scores (E-scores) were calculated relative to untreated control cells. A positive E-score denotes enrichment of the motif relative to the center of the ATAC-seq peak while a negative E-score denotes depletion of the motif. Motifs with a p_adj_ ≤ 1E-6 are colored blue. (B) E-scores for ZnCl_2_-treated cells vs TPA-treated cells show a strong negative correlation. Colors denote TF motifs that were significant (p_adj_ ≤ 1E- 6) in both conditions, one condition only, or neither condition. (C) Venn diagrams denoting the number of TF motifs that were significantly enriched or depleted in each treatment. Overlap indicates the TF motifs whose enrichments were inverted between treatments. (D) Left, DAVID tools analysis showing that motifs with a significant positive enrichment in ZnCl_2_ and a significant depletion in TPA are enriched for Zn^2+^ fingers relative to the background (all 880 TF motifs). Listed annotations are the top 15 annotations with a FDR < 0.1. Right, annotations with a FDR < 0.1 for TF motifs that were enriched with TPA and depleted in ZnCl_2_.

Many of the TF motifs that emerge from TFEA correspond to TFs that either directly bind Zn^2+^ or are known to be correlated with metal homeostasis. Functional annotation analysis using DAVID^42,43^ showed that of the TF motifs that are enriched in regions of open chromatin with ZnCl_2_ and depleted with TPA, most are zinc-finger proteins (**Figure 4D**), suggesting that the functionality of zinc-finger TFs may be sensitive to increases in the labile Zn^2+^ pool in the nM range. On the other hand, functional annotation of TF motifs that are enriched in open chromatin in TPA and depleted in ZnCl_2_ did not reveal any meaningful trends (**Figure 4D**), suggesting that perhaps these conditions lead to non-specific global changes in chromatin accessibility. For the motifs that are enriched in elevated Zn^2+^ and depleted in Zn^2+^ deficiency, many correspond to poorly annotated Zn^2+^ finger proteins, such as ZNFs and ZSCANs. However, there were some hits that are well characterized and that were particularly intriguing. For example, the CCCTC-binding factor (CTCF) is an 11-zinc finger protein involved in chromatin organization. Two CTCF motifs were globally depleted in TPA (E-score = -0.28, p_adj_ = 1E-437) and enriched in elevated Zn^2+^ (E-score = 0.42, p_adj_ = 1E-999). Our lab recently used single molecule microscopy to show that CTCF senses low Zn^2+^ conditions and becomes significantly more mobile when Zn^2+^ is depleted, suggesting decreased association with chromatin.^44^ One of the two MTF1 motifs in our database was found to be significantly enriched in elevated Zn^2+^ (E-score = 0.148, p_adj_ = 1E-14). This correlates well with our differential accessibility results showing that regions around *MT1E* and *MT2A* become more accessible with ZnCl_2_ treatment. We also observe that a motif of the zinc-finger DNA methyltransferase DNMT1 is strongly enriched in elevated Zn^2+^ (E-score = 0.47, p_adj_ = 1E-171) and depleted in TPA (E-score = -0.38, p_adj_ = 1E-278). DNMT1 is not a canonical TF but methylates DNA during DNA synthesis, and our lab has previously shown that DNA synthesis is impaired in Zn^2+^ deficiency.^33^ Finally, we find that multiple motifs associated with ELK1, a terminal TF of the MAPK pathway, show enrichment in elevated Zn^2+^ (E-score ∼ 0.25, p_adj_ ∼ 1E-30). While Elk1 is not a zinc-finger TF, previous work has shown that elevated Zn^2+^ activates the MAPK pathway via Ras GTPase, leading to phosphorylation of an Elk1-based biosensor and increases expression of the Elk1 target *FOSL1* mRNA.^37^ In summary, we find that the majority of the hits with high E-scores are enriched in regions of open chromatin upon elevation of Zn^2+^ and depleted in decreased Zn^2+^, and these largely correspond to zinc-finger TFs.

While TFEA reveals TF motifs that become enriched in newly opened chromatin upon a given treatment, it does not directly reveal whether TF occupancy on DNA has changed. To probe how TFEA enrichments correlate with TF occupancy, we selected a candidate TF and used chromatin immunoprecipitation to pull down regions of DNA bound to the TF, followed by quantitative PCR (ChIP-qPCR) to quantify how much of the TF is bound to a given target gene. As a TF target, we selected TP53, as it emerged as a strong hit in TFEA and there are multiple ChIP/ChIP-seq datasets indicating robust vetting of the antibody for ChIP. TFEA indicated that treatment with TPA enriched the TP53 motifs in open chromatin (E-score = 0.071, p_adj_ = 1.00E- 11), while addition of ZnCl_2_ depleted the TP53 motif (E-score = -0.074, p_adj_ = 1.00E-11) (**Figure 5A, Supplementary dataset 4**). TP53 contains a DNA binding domain that coordinates a single Zn^2+^ ion using three cysteine residues (C176, C238, and C242) and one histidine residue (H179).^22^ It is well established that loss of Zn^2+^, typically through the tumorigenic mutation R175H, results in misfolding and inactivation of TP53, indicating that Zn^2+^ binding is essential for TP53 activity.^22,23,45^

**Figure 5.**
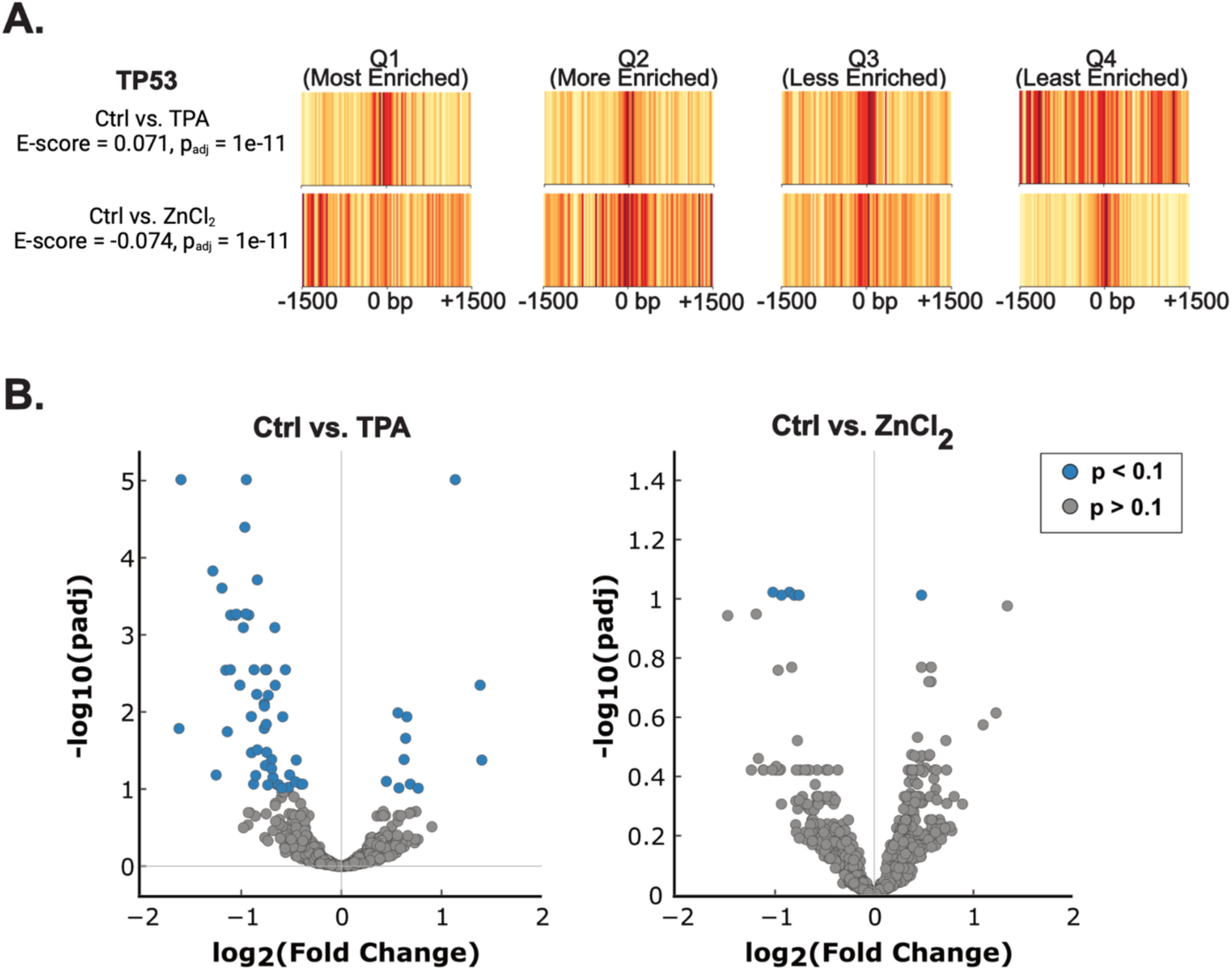
TP53 motifs show differential enrichment upon Zn^2+^ perturbation. (A) Barcode plots that show enrichment of the *TP53* motif upon treatment of MCF10A cells with TPA (top). Addition of exogenous ZnCl_2_ results in depletion of the same motif (bottom). The barcode plots represent each quartile (Q1 – Q4) of the enrichment curves generated for the TP53 motif. Red indicates more enrichment of the motif; yellow is less enrichment of the motif. (B) A subset of p53 binding sites from a ChIP-seq dataset (GSM3378513) are differentially accessible depending on cellular Zn^2+^ status. ATAC-seq reads were mapped to 2164 ChIP-seq peaks and subjected to differential accessibility analysis using DESeq2. With TPA treatment, 62 peaks showed significant (p_adj_ ≤ 0.1) changes in accessibility (51 decreased, 11 increased). With ZnCl_2_ treatment, 7 peaks showed significant changes in accessibility (1 increased, 5 decreased).

Because ATAC-seq identified more than 50,000 peaks, we sought to narrow the number of possible TP53 binding sites against which to perform ChIP-qPCR. Hence, we used a previously published TP53 ChIP-seq dataset (GSM3378513)^46^ where MCF10A cells were treated with 5 µM Nutlin-3A for 6 hours to activate TP53. We reasoned that this ChIP-seq dataset would allow us to identify bona fide regions of the genome to which TP53 binds. Further, we intersected the ChIP-seq dataset with a PRO-seq dataset (GSE227931) where MCF10A cells were treated with Nutlin-3A (10 µM, 3 hours) to identify regions of active transcription marked by the presence of bidirectional transcripts.^47^ This reduced the number of potential TP53 binding sites from approximately 13,500 (as identified in the ChIP-seq dataset) to 2164 (ChIP-seq intersected with PRO-seq). We then took this list and queried whether any of the validated TP53 binding sites were differentially accessible in response to Zn^2+^ in our ATAC-seq dataset. Differential accessibility analysis using DESeq2 revealed that with TPA treatment, 62 sites were differentially accessible (p_adj_ ≤ 0.1), with 51 of these being less accessible and 11 of these becoming more accessible (**Figure 5B**, **Supplementary Table 1**). Conversely, treatment with ZnCl_2_ resulted in 6 regions being differentially accessible, with 5 of these being less accessible and 1 being more accessible (**Figure 5B**, **Supplementary Table 2**). No sites with significant changes in accessibility were shared between both treatments.

From this list of differentially accessible sites, we selected six TP53 binding sites against which to perform ChIP-qPCR. As shown in **Figure 6**, the six target sites were selected because they show differential chromatin accessibility via ATAC-seq for at least one of the Zn^2+^ perturbations and were either within an intron or within 1000 bp of a target gene (*ERGIC1*, *NFIB*, *SFN*, *EGR1*, *PLD5*, *LRIG3-DT*). All six are associated with active transcription in MCF10A cells in response to Nutlin-3A based on PRO-seq bidirectional tracks (**Figure 6**). We also included the well-established p53 target gene *CDKN1A* (p21) as a positive control for immunoprecipitation and a negative control primer set that would not be expected to show enrichment with and without the TP53 antibody.

**Figure 6.**
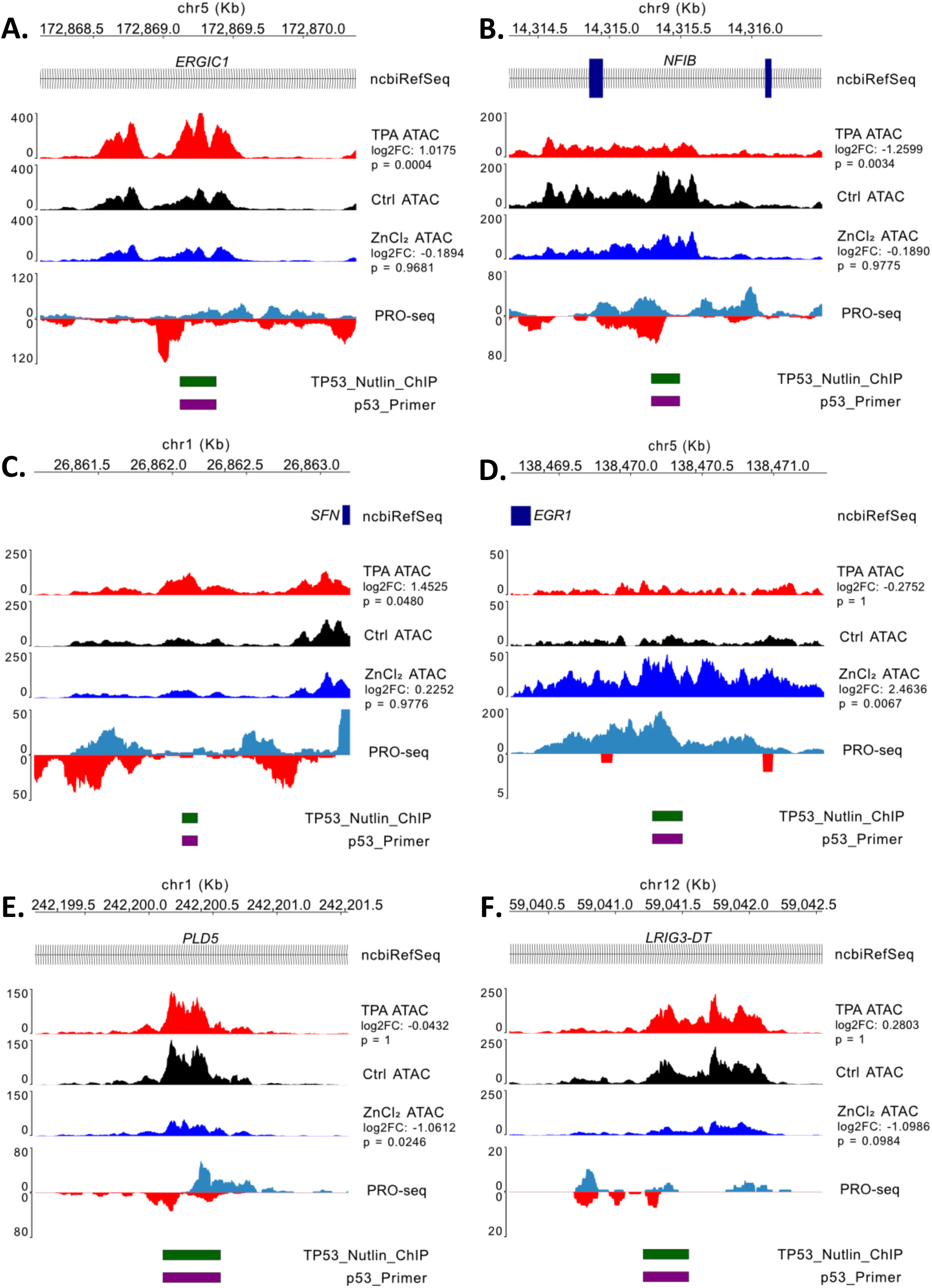
Selected regions of interest for ChIP-qPCR assays with ATAC-seq coverage tracks for cells treated with either 50 µM TPA, a media-only Control (Ctrl), or 30 µM ZnCl_2_. Also shown are the PRO-seq coverage tracks for MCF10A cells treated with 10 µM Nutlin-3A for 3 hours (GSE227931), the annotated region from the GSM3378513 p53 Nutlin-3A ChIP-seq dataset, and the predicted amplicon from ChIP-qPCR. Coverage tracks as noted above for the (A) *ERGIC1*, (B) *NFIB*, (C) *SFN*, (D) *EGR1*, (E) *PLD5*, and (F) *LRIG3-DT* regions.

We performed ChIP-qPCR against the target regions using a p53 specific antibody. As a control, we incubated chromatin with Protein A/G beads to assess the overall background signal and calculated the signal to noise ratio (SNR) by dividing the percent IP of the antibody sample by the beads-only sample (**Figure 7A**). Of our qPCR targets, *CDKN1A* (p21) had the strongest overall signal as well as SNR, as expected given that it is a well-studied TP53 target. The negative control showed no difference between the plus and minus antibody condition, further validating the ChIP conditions. Most targets had comparable signal and SNR across conditions (average SNR > 5), with control samples often displaying the largest SNR value except for *SFN* and *LRIG3-DT*, which had larger SNR values in the ZnCl_2_ samples. *EGR1* showed the lowest overall SNR across all conditions, with the TPA and control conditions displaying an average value lower than 5 (TPA: 3.62; control: 4.08; Zn: 10.80).

**Figure 7.**
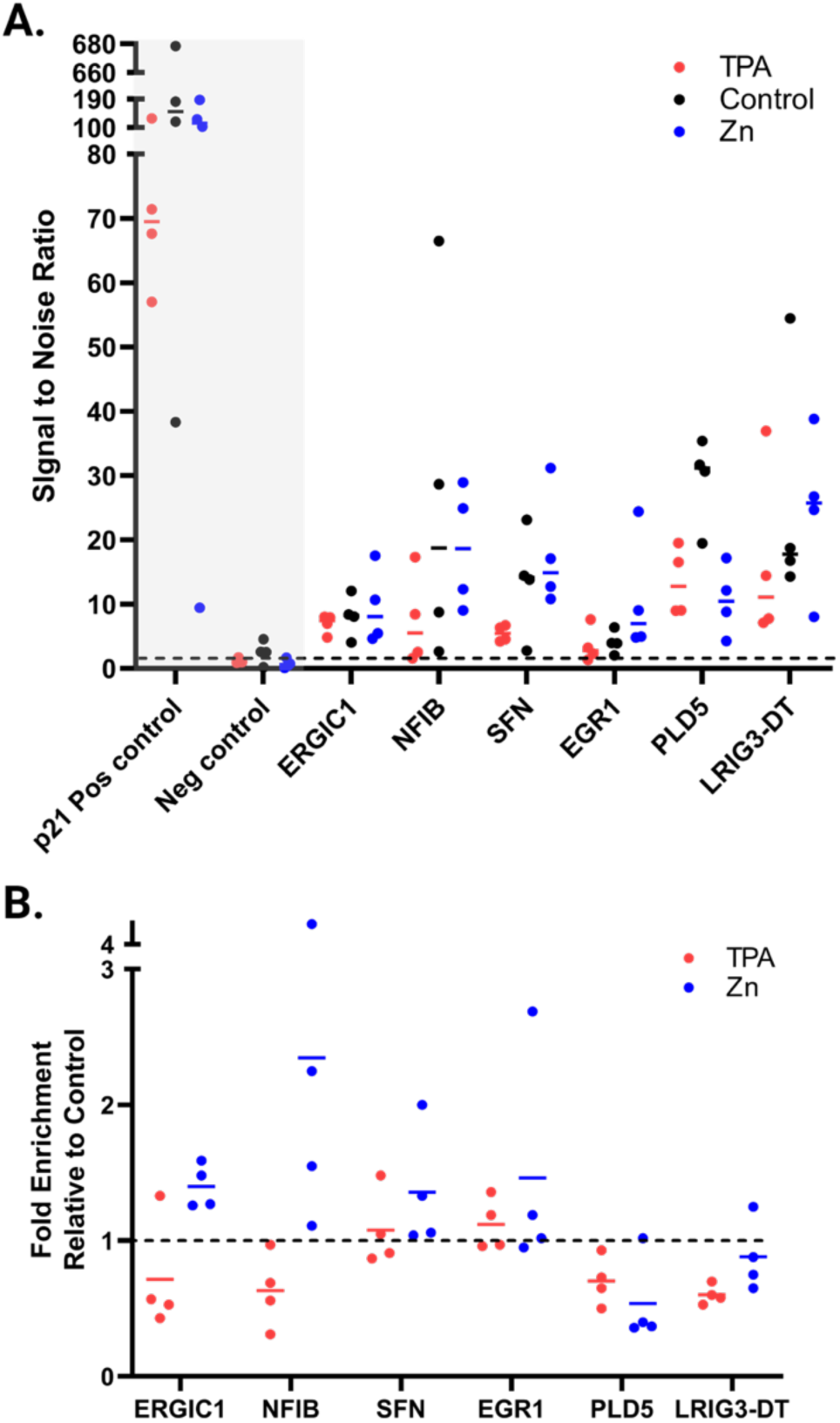
ChIP-qPCR results for putative p53 targets. (A) Signal to noise (SNR) ratios for each ChIP-qPCR target. SNR was calculated as the signal from the +p53 ChIP divided by the bead-only sample. Each data point is a biological replicate that was averaged between two technical replicates; lines within each dataset represent the mean. Dotted line denotes a SNR value of 1.5. The positive and negative controls for p53 binding are shown in the gray box. (B) Relative enrichment of each ChIP-qPCR target. Enrichment was calculated by dividing the percent pulldown (relative to input chromatin) of the treatment by the percent pulldown of the control (untreated cells). Dashed line indicates enrichment ratio of 1 (i.e. no enrichment).

We then calculated a relative fold enrichment of TP53 binding in our TPA and ZnCl_2_ treatments by dividing the treatment percent IP by the control percent IP (of samples treated with p53 antibody). The selected gene targets showed changes in TP53 binding as a consequence of Zn^2+^ perturbation, where the fold enrichment values (indicative of increased or decreased binding) largely agreed with their respective changes in accessibility (for accessibility changes where p_adj_ < 0.1) (**Figure 7B**). The TP53 binding site at the intron of *NFIB* showed a marked decrease in apparent accessibility upon treatment with TPA (**Figure 6B**) and showed a decrease in TP53 occupancy (0.63 vs. 1 for TPA vs. control). Similarly, *SFN* showed an increase in accessibility in TPA (**Figure 6C**) and its binding by TP53 was slightly increased (1.08 vs. 1 for TPA vs control). *EGR1* displayed a large positive log2foldchange in accessibility upon ZnCl_2_ treatment (**Figure 6D**) and its occupancy by TP53 was enriched (1.46 vs. 1 for Zn^2+^ vs control). Both *PLD5* and *LRIG3-DT* exhibited a decrease in accessibility when cells were treated with ZnCl_2_ (**Figures 6E-F**) and their binding by TP53 was indeed reduced (0.54 and 0.88 vs. 1, respectively, in Zn^2+^ vs control). *ERGIC1* was the one target gene where differential TP53 binding was opposite the accessibility changes by ATAC-seq. In TPA, accessibility was increased (**Figure 6A**), whereas binding by TP53 decreased (0.72 vs 1 in TPA vs control). Overall, our results indicate that Zn^2+^ induced changes in the enrichment of TF motifs in regions of chromatin accessibility are associated with changes in TF binding to target genes.

## Discussion

It has long been known that metal homeostasis is essential for proper cellular and organismal health, but relatively little has been done to probe the genomic landscape when this homeostasis is disrupted. Here, we performed ATAC-seq on MCF10A mammary epithelial cells following treatments that either raised intracellular Zn^2+^ to approximately 7 nM or depleted intracellular Zn^2+^ to less than 1 pM. In both Zn^2+^ rich and Zn^2+^ deficient states, we saw profound global changes in chromatin accessibility associated with differential enrichment of numerous TF motifs. Elevated Zn^2+^ induced changes in 449 motifs, while decreased Zn^2+^ induced changes in 322 motifs. The results indicate that changes in the labile Zn^2+^ pool alter the ability of specific TFs to access their DNA targets, with the majority of TF accessibility enriched in high Zn^2+^ and depleted in low Zn^2+^. Importantly, the Zn^2+^ perturbations used here are subtle and within the range of Zn^2+^ dynamics observed during normal physiology. For example, we recently showed that there is a pulse of labile Zn^2+^ immediately following mitosis in the mammalian cell cycle where Zn^2+^ increases to ∼ 1.6 nM for a few hours.^34^ The results presented here suggest that dynamics in labile Zn^2+^ can alter chromatin accessibility, leading to potential changes in transcriptional regulation.

Many of the differentially enriched motifs are for TFs that begin with the identifiers ZNF, ZSCAN, or ZBTB. Most of these are poorly characterized beyond the knowledge that they contain Zn^2+^ binding domains and may be involved in transcriptional regulation. However, recent work has started to explore the role of these Zn^2+^ finger TFs in the cell. One recent study found that evolutionarily conserved Zn^2+^ finger proteins were more apt to bind to promoters but Zn^2+^ finger TFs that evolved more recently are more likely to bind to transposable elements.^48^ Transposable elements have long been thought of as invaders that cause genomic instability and disease^49,50^ but more recently they have been recharacterized as potential hubs of transcriptional regulation, as on average 20% of all binding sites in TF ChIP-seq datasets colocalize with transposable elements.^51^ It is therefore possible that Zn^2+^ finger TFs regulate these sites, and it is most likely in a repressive role as work has shown that deletion of clusters of Zn^2+^ fingers in mice reactivates retrotransposons.^52^

While many of the differentially enriched motifs are for poorly characterized Zn^2+^-finger TFs, some hits are well characterized and particularly intriguing because independent studies indicate the TFs are regulated by Zn^2+^. For example, one of the motifs for MTF1 showed increased association with open chromatin in elevated Zn^2+^, which fits with a large body of work showing that in elevated Zn^2+^ MTF1 translocates to the nucleus to regulate the expression of metal homeostasis genes. Two CTCF motifs were depleted in low Zn^2+^ and enriched in elevated Zn^2+^, which is consistent with a recent study from our lab indicating that upon Zn^2+^ decrease, CTCF is more mobile with a decreased residence time on DNA, suggesting decreased association with chromatin.^44^ Finally, we found that 7 out of 8 motifs associated with ELK1, a terminal TF of the MAPK pathway, were significantly enriched in elevated Zn^2+^. While Elk1 is not a zinc-finger TF, we have shown that elevated Zn^2+^ activates the MAPK pathway, leading to phosphorylation of an Elk1-based biosensor and increased expression of the Elk1 target *FOSL1* mRNA.^37^ Combined, these results suggest that TFEA accurately predicts TFs whose activity is regulated by Zn^2+^.

Here, we used TFEA to couple differential accessibility with TF motif scanning to infer global activation or repression of TFs. Positive enrichment of a TF motif does not necessarily indicate that all the motif occurrences are bona fide TF binding sites, nor does it indicate that all TF binding sites will show an increase in accessibility. To develop a pipeline for validating TFEA predictions, we used publicly available datasets to narrow the scope. For example, TFEA applied to our ATAC-seq dataset detected > 5700 p53 motifs. We used a published p53 ChIP-seq dataset to filter these hits to genomic regions where p53 had previously been shown to bind. We further applied a p53 PRO-seq dataset to identify p53 binding sites near regions of active transcription, as identified by bidirectional transcripts. This analysis revealed only ∼2100 of the ChIP-seq binding sites showed productive transcription upon activation of p53 by a known agonist (Nutlin-3A). Of these ∼2100 binding sites, 86 showed differential accessibility in response to cellular Zn^2+^ perturbation. From this narrowed list, we selected 6 putative p53 binding sites to query with ChIP qPCR. For 5 of the 6 sites, p53 binding correlated with local accessibility, such that a decrease in accessibility led to decreased p53 binding and an increase in accessibility correlated with increased p53 binding. These results suggest that TFEA coupled with ATAC-seq is an excellent tool for profiling which TFs may be activated upon a given perturbation. With the availability of curated ChIP-seq datasets in repositories such as ENCODE and Cistrome, researchers can perform the simple and relatively inexpensive ATAC-seq, scan for enriched TF motifs using TFEA, computationally correlate changes in accessibility with candidate TF binding sites found within the literature, and then probe a subset of sites using ChIP-qPCR.

While we found that perturbing the labile Zn^2+^ pool of a cell leads to significant changes in the accessibility of many genomic sites, the mechanism leading to these chromatin accessibility changes remains an unanswered question. At any given time, there are thousands of zinc-coordinating TFs in the nucleus, whether bound to a genomic sequence or freely diffusing. Given that many of the motifs enriched in elevated Zn^2+^ and depleted in low Zn^2+^ correspond to zinc-finger TFs, one possibility is that the zinc-occupancy, and hence DNA-binding ability, of these TFs is directly influenced by the labile Zn^2+^ pool. Consistent with this possibility, we previously found that low Zn^2+^ causes increased mobility and decreased dwell-time of CTCF in the nucleus, suggesting that in low Zn^2+^ CTCF’s ability to interact with chromatin is decreased.^44^ Another possibility is that Zn^2+^ indirectly affects TF function via cell signaling. Consistent with this possibility, we found a motif associated with the non-zinc-finger TF Elk1 enriched in regions of open chromatin in elevated Zn^2+^. Previously our group has found that elevated Zn^2+^ leads to the activation of the MAPK pathway, and hence Elk1.^37^ Finally, many zinc-finger TFs directly modulate chromatin. Zn^2+^ finger TFs such as ZBTB33 are known to associate with methylated DNA, a mark of heterochromatin and gene repression,^53,54^ and the primary DNA methyltransferase that preserves methylation during cell division, DNMT1, has a Zn^2+^ binding domain that aids in its recognition of hemimethylated DNA.^55^ While the treatment window of our work was likely too short to alter DNA methylation patterns, it is possible that chronic disruption of Zn^2+^ homeostasis could perturb DNA methylation patterns, as was recently shown in *Arabidopsis*.^56^ As noted previously, CTCF is a chromatin organizing protein with 11 Zn^2+^ finger motifs that plays an important role in insulating topologically associated domains (TADs) of chromatin.^57^ Changes to the labile Zn^2+^ pool that affect CTCF’s function could disrupt TADs, as it was shown that CTCF degradation in mouse embryonic stem cells distorts TAD architecture,^58^ rendering potential changes to chromatin accessibility. Given the diversity of zinc-binding proteins, it is likely multiple mechanisms contribute to the observations in this work. In conclusion, this manuscript reveals for the first time that chromatin accessibility and transcription factor binding to DNA can be modulated by subtle changes in the labile zinc pool, suggesting that zinc may serve as a previously unrecognized modulator of transcriptional activity.

## Methods

### Molecular cloning

PiggyBac-NLS-ZapCV2 was generated by digesting PiggyBac-NES-ZapCV2^33^ with EcoRI and SalI. NLS-ZapCV2 was amplified from pcDNA(3.1)-NLS-ZapCV2 with EcoRI (5’ end) and SalI (3’ end) restriction sites using the following primers: ggaattGAATTCGCTTGGTACCGAGCATGCC and gaagcgGTCGAGCCACTGTGCTGGATATCTGCAGAA TTC. The NLS-ZapCV2 insert and PiggyBac-NES-ZapCV2^33^ were subsequently digested with EcoRI and SalI, and the NLS-ZapCV2 insert was ligated into the PiggyBac backbone.

### Cell Culture

Wildtype MCF10A cells (ATCC #CRL-10317) were cultured at 5% CO_2_ in DMEM/F12 (ThermoFisher, #11320033) supplemented with 5% Horse Serum (ThermoFisher, #16050122), 1% penicillin/streptomycin (ThermoFisher, #15070063), 20 ng/mL EGF (ThermoFisher, #), 0.5 mg/mL hydrocortisone (SigmaAldrich, #H0888), 100 ng/mL Cholera toxin (SigmaAldrich, #C8052, and 10 µg/mL insulin (ThermoFisher #12585014). To generate MCF10A cells stably expressing PiggyBac-NLS-ZapCV2, approximately 1.5 million wildtype MCF10A cells were incubated with 1 µg PiggyBac-NLS-ZapCV2 and 200 ng Super PiggyBac Transposase (Systems Biosciences #PB210PA-1) and electroporated using a Neon Electroporation System (ThermoFisher). Cells were plated into antibiotic-free media (the same composition as above, minus antibiotic) for 24 hours and then underwent puromycin selection using 0.5 µg/ml puromycin for 7 days prior to experiments.

### Live cell fluorescence microscopy

FRET sensor calibrations were performed on a Nikon widefield microscope equipped with a Ti-E perfect focus system, an iXon3 EMCCD camera (Andor), mercury arc lamp, Lambda 10-3 filter changer (Sutter Instruments), and a 20x Plan Apo air objective (Nikon). Filter settings used were: CFP (434/16 excitation, 458 dichroic, 470/24 emission), YFP FRET (434/16 excitation, 458 dichroic, 535/20 emission), and YFP (495/10 excitation, 515 dichroic, 535/20 emission). Camera exposure times of 300 ms, EM multiplier of 300, and 1 MHz readout speeds were consistent across all channels.

### Quantification of nuclear Zn^2+^ levels

MCF10A cells stably expressing PiggyBac-NLS-ZapCV2 were equilibrated at 37°C, 0% CO_2_ in phosphate-free HEPES-buffered Hanks balanced salt solution (HHBSS) containing 1.26 mM CaCl_2_, 1.1 mM MgCl_2_, 5.4 mM KCl, 20 mM HEPES, and 3 g/L glucose for 30 min prior to imaging. Baseline resting FRET ratios (R_rest_) were collected for 10 min, at which point ZnCl_2_ was added to and final media concentration of 30 µM for approximately 50 min. Cells were then washed 3x in phosphate-free HHBSS to remove excess ZnCl_2_ and 50 µM of tris(2-pyridylmethyl)amine (TPA) diluted in phosphate-free HHBSS was added to chelate Zn^2+^ and determine the sensor’s minimum FRET ratio (R_min_). Cells were washed 3x in phosphate-free HHBSS (with no CaCl_2_ or MgCl_2_) to remove excess TPA, and a solution containing 119 nM buffered Zn^2+^, ^59^ 1.5 µM pyrithione, and 0.001% w/v saponin was then added to determine the maximum FRET ratio of the sensor (R_max_). Images were then processed in MATLAB R2017b using previously published analysis code.^33,37^ Briefly, nuclei were computationally detected using the fluorescence signal in the YFP FRET channel and the background corrected FRET ratio was calculated by dividing the background corrected acceptor (YFP FRET) signal by the background corrected donor (CFP) signal. For each cell, average labile Zn^2+^ concentrations were calculated using the equation 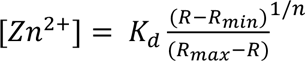 with a *K*_d_ = 5.3 nM and Hill coefficient (n) = 0.29.

### ATAC-seq sample preparation and sequencing

Approximately 50,000 MCF10A cells were pre-incubated in phosphate-free HHBSS for 30 min at 37°C and 0% CO_2_, followed by treatment with either 50 µM TPA, 30 µM ZnCl_2_, or phosphate-free HHBSS for 30 min. Nuclei isolation and transposition followed the OMNI ATAC-seq protocol as previously reported^11^ using the Illumina Tagment DNA TDE1 Enzyme & Buffer Kit (Illumina #20034197). Libraries were generated using Illumina Nextera CD Indexes (Illumina #20018708) and sequenced in paired-end mode (2×75bp) on an Illumina Nextseq 500 at the BioFrontiers Next Generation Sequencing Facility (CU Boulder).

### ATAC-seq data processing

Quality control of demultiplexed fastq files was performed using fastQC (v0.11.8).^60^ Adapters were trimmed using BBDuk^61^ using the following flags: ktrim=r, qtrim=10, k=23, mink=11, hdist=1, maq=10, minlen=20, tpe, tbo. Resulting trimmed fastq files were mapped in a paired-end fashion with HISAT2^62^ (v2.1.0) to hg38. Resulting BAM files were indexed with samtools (v1.8).^63^ Regions of open chromatin were annotated using HMMRATAC^39^; for downstream analyses (e.g. differential accessibility, TFEA, etc.), the entire open region, rather than the annotated summit, was used as an input. For assessing which regions of open chromatin were differentially accessible, peaks from all treatments were merged using muMerge.^17^ Peaks that overlapped with blacklisted genomic regions^64^ were removed using bedtools (v2.28.0). Mapped read counts were then counted using featureCounts in R (v4.0.5) and differential accessibility was determined using DESeq2.^18^

For assessment of differential TF motif enrichment using TFEA,^17^ regions of interest (ROIs) were defined by merging HMMRATAC-called peaks from two biological replicates using muMerge. Read coverages within each ROI were then calculated and differentially accessible ROIs were found using DESeq2, with ranking determined by the ROIs’ p-values. TF motif scanning was then performed within the +/- 1500 bp window surrounding each ROI center using FIMO^41^ at a p-value cutoff of 10^-5^ and a curated set of TF position weight matrices.^21^ Motif enrichment scores for each treatment relative to the control were then computed for each motif.

### PRO-seq and Library Preparation Methods

MCF10a cells were grown up in DMEM/F12 media with supplements as described in Levandowski et al.^65^ to a confluency of 70% before treatment with Nutlin-3a (10 μM Nutlin-3a in DMSO Selleck, #S8059) or vehicle (0.1% DMSO) for 3hr before the nuclei were harvested. 10×10^6^ nuclei were prepared for nuclear run-on and library preparation as described in Kwak et al.^66^ with the modifications described in Fant et al.^67^ Samples were sequenced on the Illumina NextSeq 500 platform (single-end x 75 bp). This data is publicly available on NCBI GEO at GSE227931.

### Selection of p53 binding sites for ChIP-qPCR

To find putative p53 binding sites to probe using ChIP-qPCR, we downloaded the BED file containing ChIP-seq peaks from GSM3378513^46^ (TP53 ChIP-seq in MCF10A cells treated with Nutlin-3a) from the Cistrome Data Browser (CistromeDB ID #105311). CistromeDB assesses ChIP-seq datasets on six quality control metrics (sequence quality, mapping quality, library complexity, ChIP enrichment, fraction of reads in peaks, and overlap with DNase hypersensitive sites), and this dataset passed all QC metrics; therefore, we used the BED file available from the CistromeDB website. To increase confidence in p53 ChiP-seq peaks that correlate with regions of productive transcription, we used bedtools intersect with the flag “-F 0.5” to only retain ChIP peaks that overlapped at least 50% with regions of productive transcription. This reduced the number of p53 ChIP peaks from 13,453 to 2163. We then used the featureCounts module in R to count the number of ATAC-seq reads mapped to these peaks and performed differential accessibility analysis using DESeq2. The list of significant (p_adj_ ≤ 0.1) hits was then manually curated to select for regions that were proximal to or within genes undergoing bidirectional transcription using the Nutlin-3A PRO-seq dataset. Of these, we selected six candidates whose target genes were of interest and lent themselves to successful qPCR primer validation via amplification efficiency testing.

The six candidates were selected based on the following criteria: 1) they exhibited differential accessibility in at least one of the Zn^2+^ perturbations, 2) they overlapped a p53 binding site identified by ChIP-seq, 3) they correlated with regions of active transcription upon activation of p53 by nutlin 3a, 4) primers could be designed within the introns of the gene or at most ∼1000 base pairs up/downstream of the gene. Within the p53 targets in this list is an intronic region of endoplasmic reticulum-Golgi intermediate compartment 1 (*ERGIC1*), which encodes a cycling membrane protein whose potential function is to transport cargo from the ER to the Golgi.^68^ In TPA treated cells, the accessibility at this genomic region was slightly, but significantly increased, while treatment with ZnCl_2_ did not significantly alter its accessibility. Additionally, an intronic region of the Nuclear Factor I B (*NFIB*), a TF that regulates differentiation and proliferation of the central nervous system,^69^ showed significantly reduced accessibility with TPA. We also targeted a p53 binding site upstream of stratifin (*SFN*), which, when activated by p53 because of detected DNA damage, leads to cell cycle arrest at the G2/M transition.^70,71^ This region saw a significant increase in accessibility in TPA. A region that saw a large increase in accessibility under ZnCl_2_ conditions was a p53 binding site downstream of the Early growth response 1 (*EGR1*) gene, which encodes a TF containing three C2H2-type zinc-fingers that can regulate p53 activity upon ionizing radiation exposure.^72^ Cells treated with ZnCl_2_ also displayed a slight but significant reduction in the accessibility at the intronic regions of *PLD5* and *LRIG3-DT*. Phospholipase D family member 5 (*PLD5*) has been shown to be regulated by a tumor suppressor microRNA that is itself a client of p53^73^; while the leucine-rich repeats and immunoglobulin-like domains divergent transcript (*LRIG3-DT*) encodes an integral plasma membrane protein whose expression has been shown to positively correlate with the expression of p53 in cervical intraepithelial neoplasia.^74^

As a positive control for immunoprecipitation, we used previously published^75^ primers targeting the p53 response element within the promoter of *CDKN1A* (*p21*), a well-established p53 target.^76,77^ For *CDKN1A*, the p53 binding site was not included in our DESeq2 analysis, as the primary region of active transcription within the p53 PRO-seq dataset was downstream of the p53 binding site we chose. Visually, however, the region appeared to be slightly more accessible in TPA. As a negative control for immunoprecipitation, the Human Negative Control Primer Set 1 (ActiveMotif, #71001) amplifies a 78 base pair fragment from a gene desert on human chromosome 12 and would not be expected to show enrichment when incubated with the p53 antibody.

### ChIP-qPCR

Wildtype MCF10A cells (2x 15 cm dishes with approximately 12 million cells/dish) were treated as noted above for ATAC-seq for 30 min, followed by washing with room temperature PBS, crosslinking using 1% methanol-free formaldehyde (Thermosphere #28908) for 10 min and quenching for 8 min using 125 mM glycine. Cells were washed using cold PBS and nuclei were isolated using a hypotonic buffer (10 mM Tris-HCl pH 7.5, 50 mM NaCl, 2 mM EDTA, 1% NP-40, 1 mM DTT, and 1x protease inhibitors (ThermoFisher #78441)). Nuclei were then lysed and subjected to chromatin fragmentation using a BioRuptor (Diagenode) sonicator (7×10 minute pulses, high setting) resulting in a mean chromatin length of ∼200 bp, followed by a spin at 21,130x g for 15 min at 4°C to pellet insoluble material. Chromatin was aliquoted, flash frozen in liquid nitrogen, and stored at -70°C.

To immunoprecipitate p53-bound chromatin, 200 µL of chromatin (from approximately 7×10^6^ MCF10A cells) were pre-cleared per condition by incubating with Protein A/G beads (Santa Cruz #SC-2003. Lot#B2323) for two hours at 4°C on a rotator. Then, 1/10^th^ of the pre-cleared chromatin was saved as input and frozen, and half of the remaining chromatin was rotated overnight at 4°C with 4.5 µg of anti-p53 (BD Biosciences #554293, Lot #9046678). The other half of the remaining chromatin was also rotated overnight as a no-antibody control. The following morning, chromatin samples (+/- antibody) were incubated for two hours with Protein A/G beads (15 µL of beads per 100 µL of chromatin) that had been blocked overnight with 0.5 mg/mL bovine serum albumin and 0.4 mg/mL yeast RNA. Beads were then washed twice with each a low salt buffer (20 mM Tris-HCl pH 8, 150 mM NaCl, 2 mM EDTA, 1% Triton-X 100, 0.1% SDS), a high salt buffer (20 mM Tris-HCl pH 8, 500 mM NaCl, 2 mM EDTA, 1% Triton X-100, 0.1% SDS), a high salt buffer containing LiCl (20 mM Tris-HCl pH 8, 250 mM LiCl, 1. mM EDTA, 1% sodium deoxycholate, 1% NP-40), and finally with a TE buffer (10 mM Tris-HCl pH 8, 1 mM EDTA). Chromatin samples and inputs were eluted in 200 µL of elution buffer (50 mM Tris-HCl pH 8, 10 mM EDTA, 1% SDS) for 1 hour at 37°C with shaking. To reverse crosslinks, samples and inputs were incubated overnight in 200 mM NaCl at 65°C. The following morning, all samples were diluted 1:1 with Milli Q water to lower the salt concentration and proteins were digested using 160 μg Proteinase K at 55°C for 1 hour. DNA was purified via phenol:chloroform:isoamyl alcohol (ThermoFisher #15593031) extraction and ethanol precipitation. DNA pellets from the +/- antibody samples and from the inputs were then resuspended in 27 µL or 100 µL of diethylpyrocarbonate (DEPC)-treated water, respectively.

For qPCR, 10-fold serial dilutions (1:10, 1:100, 1:1000) of each input were prepared for each primer set. qPCR reactions were prepared as follows: 600 nM forward primer, 600 nM reverse primer, 3 µL of DNA (sample or diluted input), and 1X SYBR Select qPCR master mix (ThermoFisher #4472918), with a final volume of 21 µL. qPCR was then performed in technical duplicate (10 µL/reaction) on a BioRad C1000 Touch thermal cycler using the recommended SYBR Select protocol and an annealing temperature of 57°C. C_T_ values between technical replicates were averaged and the percent IP for each treatment and primer set was calculated by aligning and comparing the signal from treatment wells to the standard curve of input chromatin given the equation: %*IP* = (*DNA quantity*) ∗ 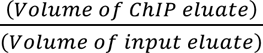 ∗ 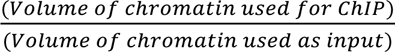. All qPCR primers used in this study are listed in **Supplementary Table 3**. The entirety of the ChIP qPCR protocol was run on 4 biological replicates.

## Data Access

All raw and processed sequencing data generated in this study have been submitted to the NCBI Gene Expression Omnibus (GEO; https://www.ncbi.nlm.nih.gov/geo/) under accession number GSE212763.

## Acknowledgements

We would like to thank the following sources for financial support: NIGMS MIRA R35 GM139644 (AEP), NIH/CU Molecular Biophysics Program and NIH Biophysics Training Grant T32 (GM-065103 for LJD, GM-145437 for DO). NIH R01GM125871 (RDD, MAA), NSF ABI 1759949 (RDD, LS), and NIH R03AG061466 (RDD, TJ). We would like to thank Dr. Joe Dragavon of the BioFrontiers Institute Advanced Light Microscopy Core for assistance with microscopy and Dr. Amber Scott, Dr. Joe Cardiello, and Kevyn Jackson of the BioFrontiers Institute Next Generation Sequencing Facility for assisting with sequencing data collection. We would also like to acknowledge the University of Colorado Biochemistry Cell Culture Core Facility, especially Dr. Theresa Nahreini, for providing resources and support for all our cell work. We thank Dr. Roy Parker and Dr. Thomas Cech for the use of their laboratories’ equipment. We would also like to thank members of the Dowell & Allen lab, especially Dr. Gilson Sanchez, for fruitful discussions about chromatin accessibility. We thank Dr. Jennifer Kugel for advice on optimizing the signal to noise ratio of our ChIP-qPCR studies.

